# PTH decreases *in vitro* human cartilage regeneration without affecting hypertrophic differentiation

**DOI:** 10.1101/560771

**Authors:** Marijn Rutgers, Frances Bach, Luciënne Vonk, Mattie van Rijen, Vanessa Akrum, Antonette van Boxtel, Wouter Dhert, Laura Creemers

## Abstract

Regenerated cartilage formed after Autologous Chondrocyte Implantation may be of suboptimal quality due to postulated hypertrophic changes. Parathyroid hormone-related peptide, containing the parathyroid hormone sequence (PTHrP 1-34), enhances cartilage growth during development and inhibits hypertrophic differentiation of mesenchymal stromal cells (MSCs) and growth plate chondrocytes. This study aims to determine whether human articular chondrocytes respond correspondingly. Healthy human articular cartilage-derived chondrocytes (*n*=6 donors) were cultured on type II collagen-coated transwells with/without 0.1 or 1.0 μM PTH from day 0, 9, or 21 until the end of culture (day 28). Extracellular matrix production, (pre)hypertrophy and PTH signaling were assessed by RT-qPCR and/or immunohistochemistry for collagen type I, II, X, RUNX2, MMP13, PTHR1 and IHH and by determining glycosaminoglycan production and DNA content. The Bern score assessed cartilage quality by histology. Regardless of the concentration and initiation of supplementation, PTH treatment significantly decreased DNA and glycosaminoglycan content and reduced the Bern score compared with controls. Type I collagen deposition was increased, whereas PTHR1 expression and type II collagen deposition were decreased by PTH supplementation. Expression of the (pre)hypertrophic markers MMP13, RUNX2, IHH and type X collagen were not affected by PTH. In conclusion, PTH supplementation to healthy human articular chondrocytes did not affect hypertrophic differentiation, but negatively influenced cartilage quality, the tissues’ extracellular matrix and cell content. Although PTH may be an effective inhibitor of hypertrophic differentiation in MSC-based cartilage repair, care may be warranted in applying accessory PTH treatment due to its effects on articular chondrocytes.

## Introduction

Autologous chondrocyte implantation (ACI) is an effective treatment in patients with medium-sized cartilage defects (1). Chondrocytes isolated from healthy non weight-bearing cartilage and re-transplanted after *in vitro* expansion ideally fill the void with hyaline neocartilage (2). Variable results have, however, been found regarding the obtained cartilage quality, with fibrous or even hypertrophically differentiated tissue instead of healthy cartilage (3). Similarly, MSC-based regeneration either as part of microfracture procedures or as exogenous cell source has been shown to result in hypertrophic differentiation (4). A possible tool to prevent this unfavorable differentiation pathway may be the co-administration of parathyroid hormone related-peptide (PTHrP).

PTHrP plays an important role in early development and limb growth (5) and is crucial in maintaining the chondrocytic phenotype in native cartilage. In the growth plate, a highly organized cartilage structure that enables longitudinal bone growth, PTHrP maintains chondrocytes in a proliferating state and prevents hypertrophic differentiation and bone formation (6). Parathyroid hormone (PTH) is assumed to have similar effects as PTHrP (7,8), as they share their N-terminus and receptor (PTHR1) (9,10). Both PTH and PTHrP can enhance cartilage formation by stimulating the expression of SRY-box 9 (6) (*SOX9*, transcription factor required for chondrocyte differentiation and cartilage formation (11)) and by increasing cell proliferation through induction of cyclin D1 (*CCND1)* (12). PTHrP/PTH have been demonstrated to stimulate chondrogenic differentiation of mesenchymal stromal cells (MSCs) and to prevent hypertrophic differentiation of MSCs (13–17) and growth plate chondrocytes (18,19) *in vitro*. Lastly, in rabbit osteochondral defects, intra-articular PTH administration stimulated tissue regeneration *in vivo* (20,21). In contrast to osteochondral defect healing, ACI is based on chondrocyte implantation, either or not supplemented with MSCs (22). Although expanding chondrocytes have stem cell-like properties (23), multilineage (especially osteogenic and adipogenic) differentiation efficacy is very low compared with MSCs, which may indicate that a chondroid precursor, but not a multipotent mesenchymal cell type is present in expanding chondrocytes (24).

As yet, it is unknown whether PTH may have similar effects on articular chondrocyte-mediated regeneration in terms of inhibition of hypertrophy and stimulation of regeneration. Therefore, in order to define whether PTH holds promise as an additive treatment strategy to current challenges faced by ACI, this study determined the effects of PTH on expanded human articular chondrocytes in an *in vitro* model of cartilage regeneration.

## Materials and Methods

### Human chondrocytes

Healthy human femoral knee cartilage of three male and three female donors (mean age 68, range 47-83 years) was obtained *post-mortem*. Only macroscopically intact (Collins grade 0-1 (25)) cartilage was used (four samples grade 0, two samples grade 1). Collection of all patient material was done according to the Medical Ethical regulations of the University Medical Center Utrecht and according to the guideline ‘good use of redundant tissue for clinical research’ constructed by the Dutch Federation of Medical Research Societies on collection of redundant tissue for research (www.fedara.org). This study does not meet the definition of human subjects research or require informed consent. Anonymous use of redundant tissue for research purposes is part of the standard treatment agreement with patients in our hospital (26).

### Chondrocyte isolation and expansion

Cartilage was digested overnight in Dulbecco’s modified Eagle Medium (DMEM; 42430, Invitrogen) containing 0.1% w/v collagenase type II (CLS2, Worthington), 1% v/v penicillin/streptomycin (P/S; 15140, Invitrogen). Isolated chondrocytes were washed in PBS and expanded at 5000 cells/cm^2^ in DMEM containing 1% P/S, 10% v/v fetal bovine serum (DE14-801F, Lonza), 10 ng/mL FGF (223-FB, R&D Systems). At 80% confluency, cells were trypsinized and passaged. Passage two cells were used for redifferentiation culture or snap-frozen for RNA analysis.

### Redifferentiation culture

Since more recent ACI versions are based on collagen carriers (27), the chondrocytes were cultured on collagen type II-coated membranes in a 24-wells transwell system as described previously (28). Passage two chondrocytes were seeded on inserts (PICM01250, Millipore) with a hydrophilic poly-tetrafluoroethylene (PTFE) membrane at 1.6×10^6^ cells/cm^2^ in DMEM with 2% w/v ITSx (51500, Invitrogen), 2% w/v ascorbic acid (A8960, Sigma-Aldrich), 2% w/v human serum albumin (HS-440, Seracare Life Sciences), 100 units/mL penicillin, 100 μg/mL streptomycin, and 10 ng/mL TGF-β_2_ (302-B2, R&D Systems). Before culture, the membranes had been coated with 0.125 mg/mL type II collagen (C9301, Sigma-Aldrich) in 0.1 M acetic acid. After thorough rinsing, efficient coating was verified by immunohistochemistry (29). To mimic the different phases of maturation, PTH was supplied at different time points in two different concentrations: 0.1 or 1.0 μM PTH (based on Kafienah *et al.* (2007) (13)) was added every media change (three times a week) from days 0 (*n*=4), 9 (*n*=6) or 21 (*n*=6) onwards, whereas the controls (*n*=6) did not receive PTH. After 28 days, the tissues were fixed in 10% w/v neutral buffered formalin (for histological analysis) or snap-frozen and stored at −20 °C (glycosaminoglycan (GAG) and DNA content analysis) or −80 °C (for RNA analysis).

### Gene expression analysis

After separation of the neotissue from the transwell membranes with a scalpel (Swann-Morton), RLT with β-mercaptoethanol (M6250, Sigma) was added to the samples. Thereafter, samples were crushed and pestled with liquid nitrogen and 21G syringes until a homogeneous solution was obtained. RNA isolation was performed using the RNeasy Micro Kit (74004, Qiagen) according to the manufacturer’s protocol with DNAse treatment. RNA concentrations were measured using the NanoDrop® ND-1000 spectrophotometer (NanoDrop Technologies). Sufficient RNA purity was assumed with 260/280nm ratio >1.9. Subsequently, 500 ng RNA was reverse transcribed using the iScript cDNA synthesis Kit (Bio-Rad). RT-qPCR was performed according to the manufacturer’s protocol (technical duplicates) using TaqMan® Gene Expression Assays (Applied Biosystems). Primers for hypertrophic differentiation and (cartilaginous) extracellular matrix production were used: *COL1A1* (Hs00164004_m1), *COL2A1* (Hs00264051_m1), *COL10A1* (Hs00166657_m1), *RUN*X2 (Hs00231692_m1, recognizes all three isoforms), and *MMP13* (Hs00942589_m1), all corrected for housekeeping gene *ACTB* (β-actin) (Hs99999903_m1).

### Histological analysis

Histological evaluation was performed on 5 µm thick deparaffinized sections using Safranin O/Fast Green staining as described previously (30). Cartilage quality by histology was assessed with the Bern Score (31), specifically designed for engineered cartilage, assessing intensity and uniformity of Safranin O staining, amount and organization of the extracellular matrix and cell morphology. The maximum score is 9, representing healthy hyaline cartilage.

### Immunohistochemistry

Immunohistochemical staining was performed for (cartilaginous) extracellular matrix production and (pre)hypertrophic differentiation: types I, II and X collagen, RUNX2, IHH and PTHR1. The sections were blocked with 0.3% H_2_O_2_ in PBS and 5% PBS/BSA (type I, II, and X collagen) or TBS/BSA 5% (RUNX2) and incubated overnight at 4 °C with mouse monoclonal antibodies against type I collagen (20 μg/mL, CP-17, Calbiochem), type II collagen (0.4 μg/mL, ascites II-II6B3, Developmental Studies Hybridoma Bank), type X collagen (1:50 (actual concentration unknown), X53, Quartett) and RUNX2 (5 μg/mL, SC-101145, Santa Cruz) in 5% PBS/BSA. Subsequently, the samples were incubated with biotinylated sheep anti-mouse (1:200, RPN1001V, GE Healthcare) followed by peroxidase labeled streptavidin (1:500, IM0309 Beckman Coulter) (type I and X collagen) or goat anti mouse-HRP (1:100, P0447, DAKO) (type II collagen, RUNX2) for 1 hour in 5% PBS/BSA. Sections were counterstained with hematoxylin. Juvenile growth plate cartilage was used as a positive control. Sections were incubated with isotype antibodies as negative controls. Negative control sections did not show staining, whereas positive control sections showed specific staining (data not shown).

To determine IHH (sc-1196, Santa cruz, 4 μg/mL) and PTHR1 (sc-12777, Santa Cruz, 10 μg/mL) expression, sections were immunohistochemically stained using the ImmunoCruz™ goat LSAB Staining System (sc-2053, Santa Cruz) with citrate buffer (10 mM, pH 6) antigen retrieval (30 minutes, 70°C) (18).

### GAG and DNA analysis

GAG and DNA content of the tissue constructs were digested in 2% papain (P3125, Sigma-Aldrich) in 50 mM phosphate buffer, 2 mM N-acetylcysteine, and 2 mM Na_2_-EDTA (pH 6.5) at 65°C for 2 hours. GAGs were precipitated from tissue digests and culture medium (each medium change) using Alcian blue dye (Alcian blue 8GX, 05500, Sigma-Aldrich) saturated in 0.1 M sodium acetate buffer containing 0.3 M MgCl_2_ (pH 6.2) for 30 min at 37°C (32). Absorbance was quantified photospectrometrically (520 nm) with chondroitin sulphate

(C4384, Sigma) as reference. DNA was quantified by Hoechst 33258 (94403, Sigma) staining (33) followed by fluorescence measurement (Cytofluor, Bio-Rad) with calf thymus DNA (D4764, Sigma) as reference.

### Statistics

Statistical analyses were performed using IBM SPSS 20 for Windows (IBM SPSS Inc.). Data were tested for normality using the Kolmogorov–Smirnov test and for homogeneity of variances using the Levene’s equality of variances test. Subsequently, mixed model analysis (random effect: donor, fixed effect: PTH concentration and moment of initiating treatment) was performed. After performing a Bonferroni *posthoc* test for multiple comparisons, *p*<0.05 was considered statistically significant.

## Results

### PCR

In all PTH-treated neotissues, *COL2A1* mRNA expression was decreased compared with controls (*p*=0.042, Fig. 1). No effect of PTH was observed on *COL1A1*, *COL10A1*, *MMP13*, or *RUNX2* expression.

**Fig. 1.**
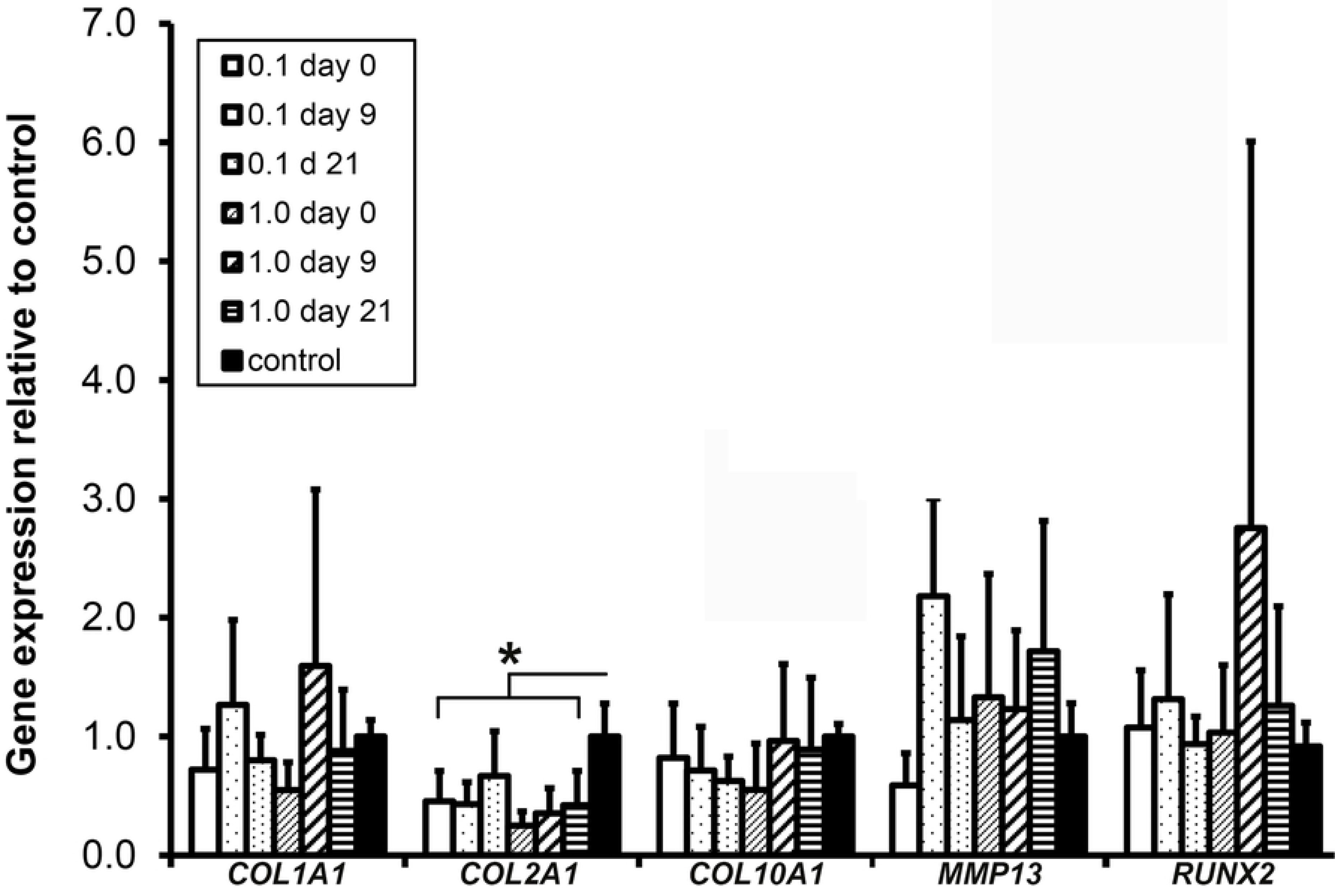
Gene expression of type I, II and X collagen (*COL1A1, COL2A1, COL10A1*), matrix metallopeptidase 13 (*MMP13*) and runt related transcription factor 2 (*RUNX2*) by human chondrocytes cultured for 28 days in the presence of 0.1 or 1.0 μM PTH from day 0, 9 or 21 onwards. Controls did not receive PTH. 0.1 or 1 µM PTH was added from day 0, 9 or 21 onwards. *: *p*=0.042. Graphs show mean ± 95% C-I. *n*=6.

### Bern score and immunohistochemistry

Both 0.1 µM PTH (*p*=0.006) and 1 µM PTH (*p*=0.022) treatment resulted in lower Bern scores compared with controls, regardless the moment of initiating treatment (Fig. 2). Safranin O staining intensity was not different between conditions. Type I collagen deposition appeared slightly higher and type II collagen deposition slightly lower in all PTH-treated cultures compared with controls (Fig. 3). Only minor type X collagen deposition was observed, with no differential expression between conditions. RUNX2 expression was present in three out of six donors, independent of PTH addition. IHH was abundantly present in all donors and conditions, but was not differentially expressed. PTHR1 expression appeared decreased by 0.1 and 1.0 μM PTH treatment (irrespective of initiation of treatment) compared with controls.

**Fig. 2.**
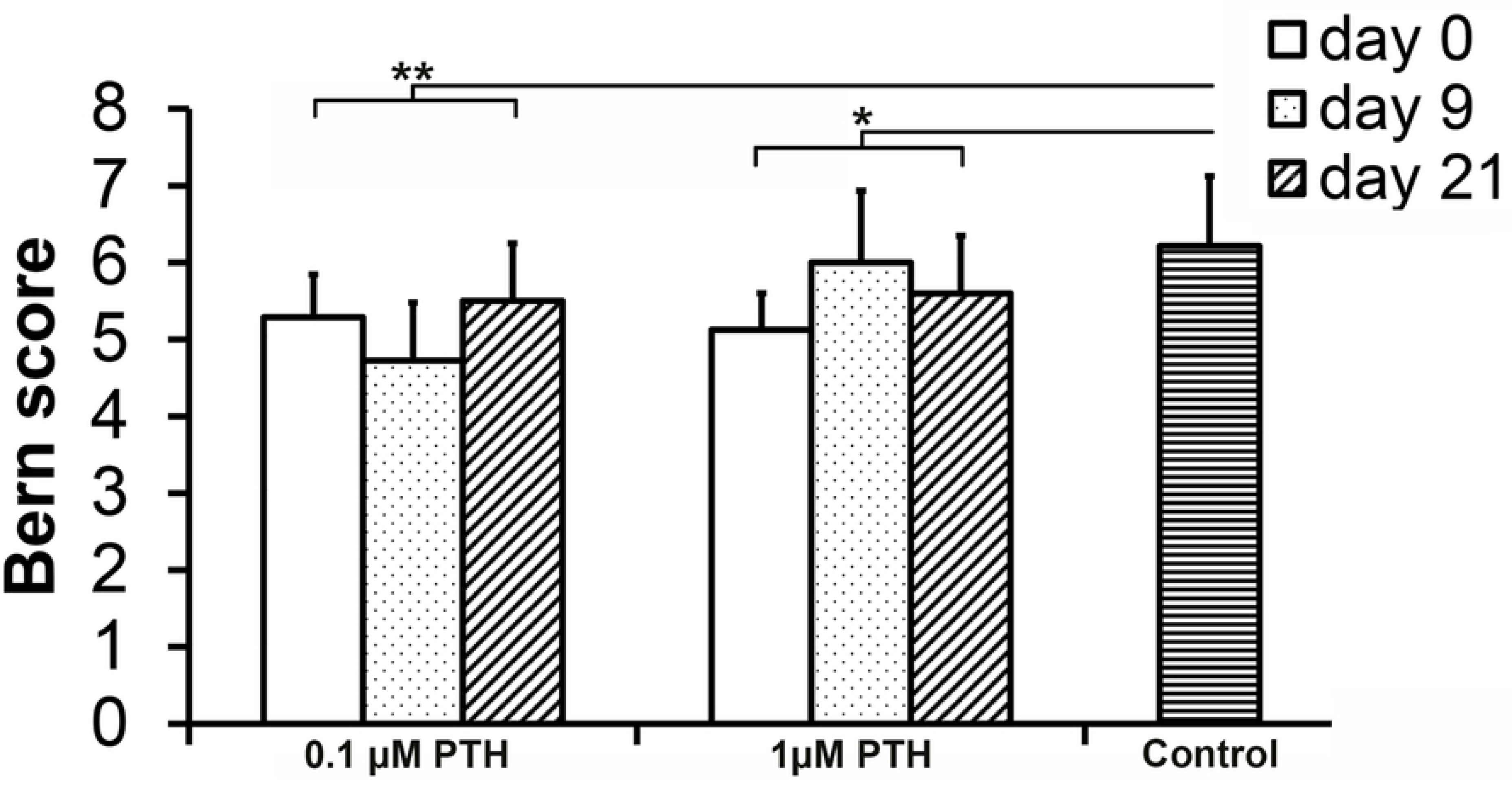
Bern score of human chondrocytes cultured for 28 days in the presence of 0.1 or 1.0 μM PTH from day 0, 9 or 21 onwards. Controls did not receive PTH. 0.1 µM and 1 µM PTH addition resulted in lower Bern scores compared with controls (*: *p*=0.006, **: *p*= 0.022, respectively). No differences were observed between PTH concentrations or moment of supplementation. Graphs show mean ± 95% C-I. *n*=6.

**Fig. 3.**
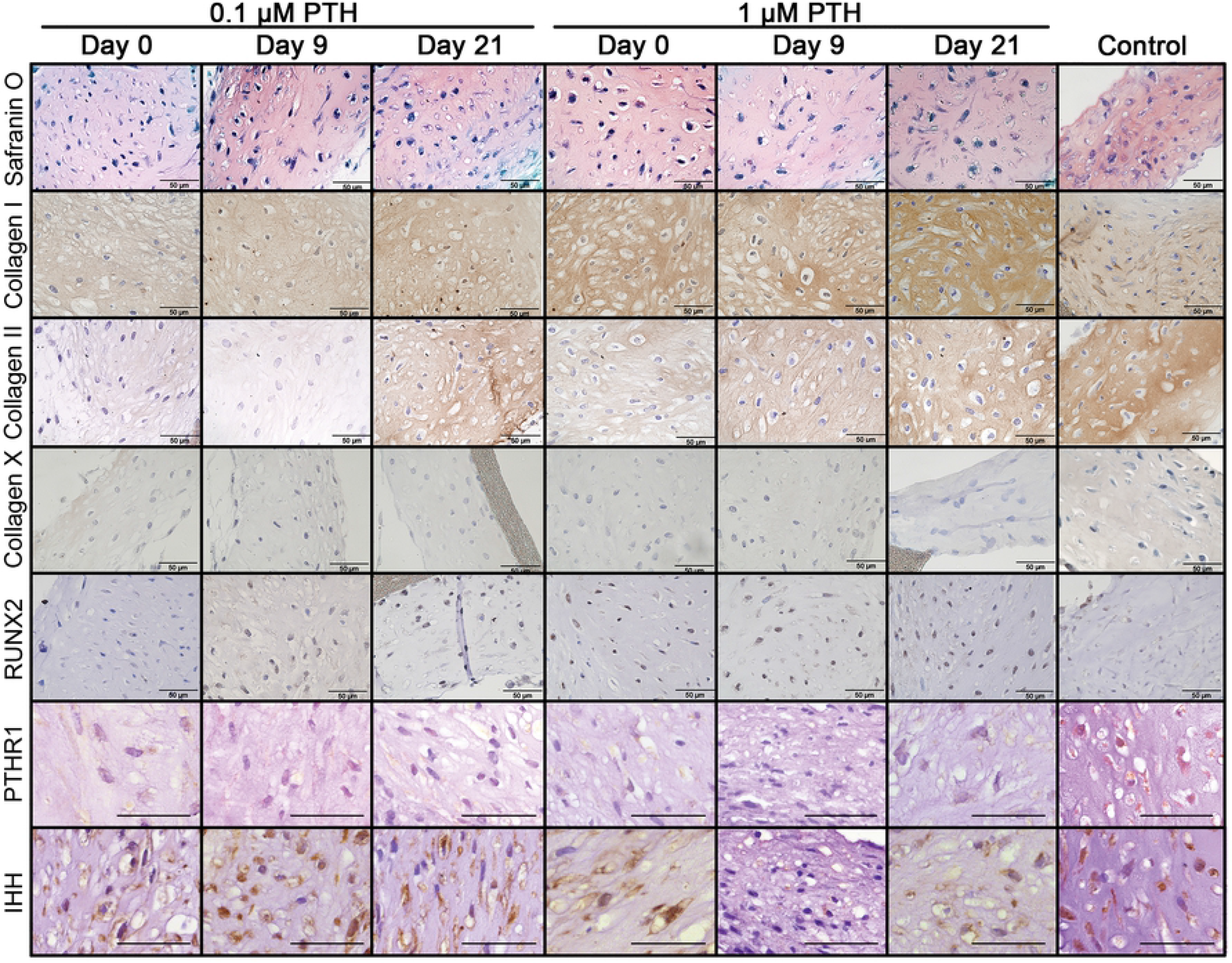
Histological sections of human chondrocytes cultured for 28 days in the presence of 0.1 or 1.0 μM PTH from day 0, 9 or 21 onwards. Controls did not receive PTH. All scale bar represent 50 µm. *n*=6.

### GAG and DNA content, GAG production and release

PTH treatment decreased the constructs’ GAG content compared with controls, regardless of the concentration or initiation of treatment (*p*=0.001, Fig. 4a). Also, DNA content of the constructs was decreased by the addition of 0.1 and 1.0 μM PTH (*p*=0.030, Fig. 4b). For both PTH concentrations, DNA content was inversely related to the duration of PTH exposure (*p*=0.027). The GAG content corrected for DNA content (indirect measure of GAG deposition per cell) was not influenced by the addition of PTH (*p*=0.482, Fig. 4c). GAG release and total GAG production (sum of total amount of GAGs released into the medium and GAG content of the regenerated tissue) were also not affected by PTH treatment (*p*=0.586, Fig. 5a,b). Relative GAG release (release as percentage of total production) did not differ between the different PTH concentrations or controls (*p*=0.656, Fig. 5c).

**Fig. 4.**
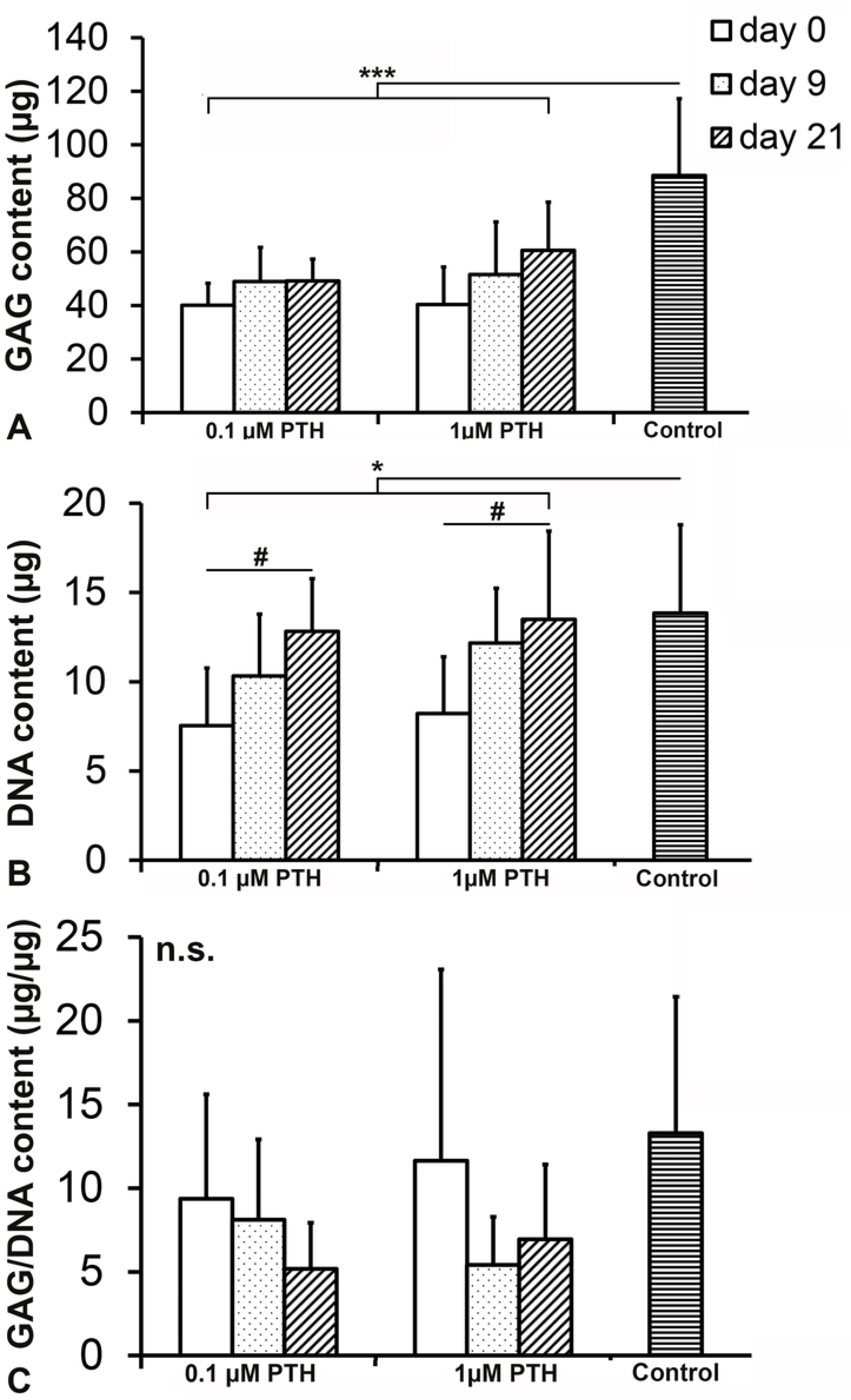
GAG, DNA and GAG/DNA content of human chondrocytes cultured for 28 days in the presence of 0.1 or 1.0 μM PTH from day 0, 9 or 21 onwards. Controls did not receive PTH. (a) PTH treatment resulted in a lower GAG content (*p*=0.001). (b) DNA content was decreased by addition of 0.1 and 1.0 μM PTH (*: *p*=0.030). A time-dependent increase in DNA content was observed in the presence of PTH (#: *p*=0.027), with no difference between 0.1 and 1.0 μM. (c) GAG/DNA content did not differ between conditions. n.s.: not significant. Graphs show mean ± 95% C-I. *n*=6.

**Fig. 5.**
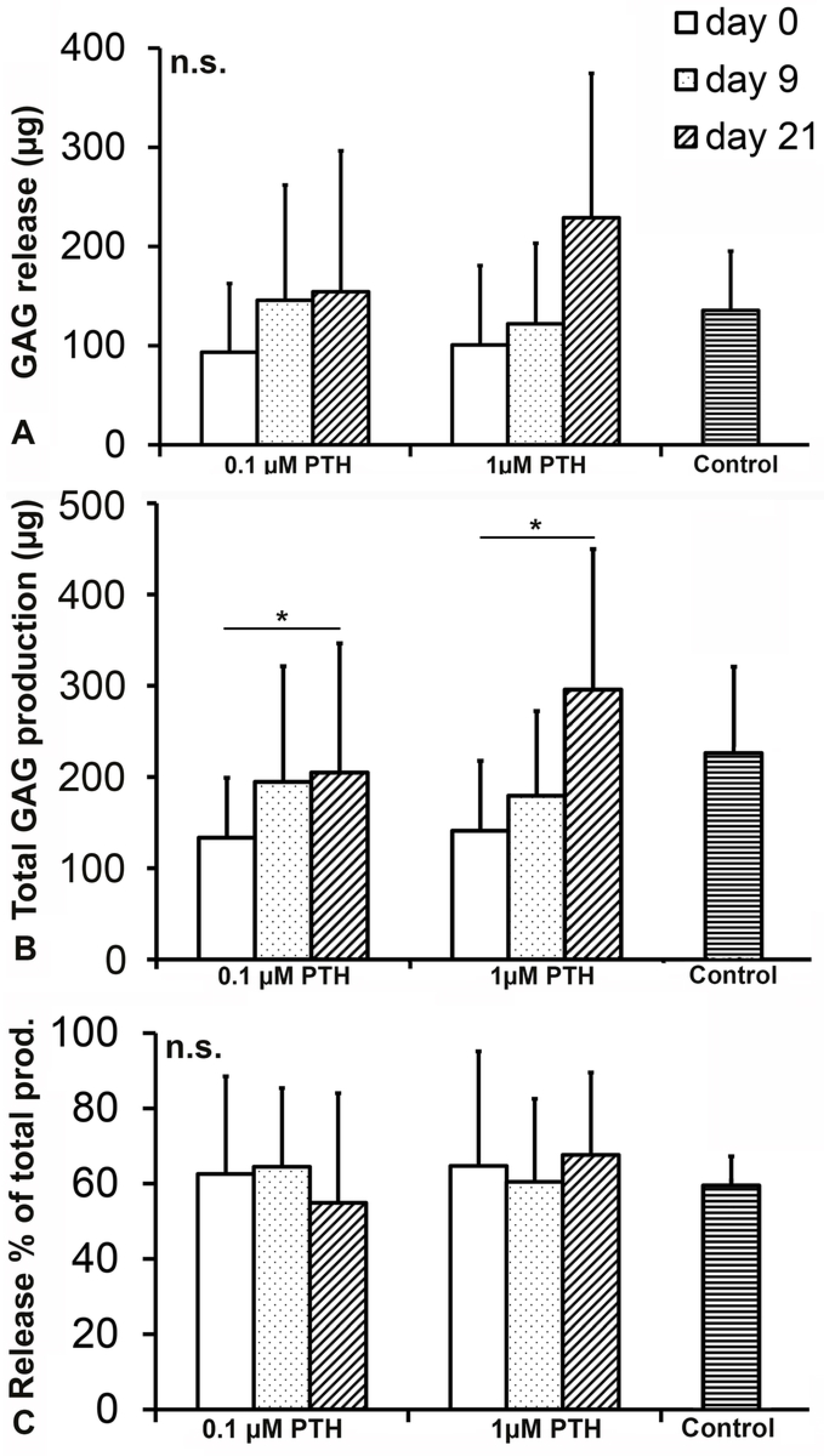
Total GAG production, total GAG release and GAG release as percentage of total production of human chondrocytes cultured for 28 days in the presence of 0.1 or 1.0 μM PTH from day 0, 9 or 21 onwards. Controls did not receive PTH. Total GAG production was dependent on moment of PTH supplementation (*: *p*=0.042), but not on PTH concentration. n.s.: not significant. Graphs show mean ± 95% C-I. *n*=6.

## Discussion

### PTH did not affect hypertrophic chondrocyte differentiation

Although ACI is effective at treating medium-sized cartilage defects(1), the newly formed cartilage can be of reduced (*e.g.* hypertrophic) quality (3). PTH and PTHrP have the ability to inhibit hypertrophic differentiation of MSCs (13–17) and growth plate chondrocytes (18,19). Therefore, the first aim of this study was to determine whether PTH could also inhibit hypertrophic differentiation in healthy human chondrocytes, the cell source used for ACI. Although both PTHR1 and IHH are considered prehypertrophic differentiation markers (34), only PTHR1 expression was inhibited by PTH supplementation, in line with previous work on growth plate chondrocytes (8,18). Since IHH, RUNX2 and type X collagen (hypertrophic differentiation marker (34)) expressions were not inhibited by PTH, the downregulation of PTHR1 presumably reflects a negative feedback mechanism to prevent overstimulation (18), rather than an inhibition of (pre)hypertrophic differentiation. The absence of a PTH-mediated effect on hypertrophic differentiation is in contrast with previously observed effects on hypertrophic articular (35) and growth plate (18) chondrocytes. Although diminished responsivity to PTH has been reported for chondrocytes (13,19), this is unlikely the cause of the lack of effect on hypertrophic differentiation, since the DNA content and associated extracellular matrix composition were affected by PTH. Presumably, the chondrocytes reached at least a prehypertrophic state in the current study, since PTHR1 and IHH were abundantly expressed in controls. However, since no clear features of hypertrophic differentiation (*e.g.* hypertrophic cellular phenotype, abundant type X collagen deposition) were observed, other factors may have been required for its induction, such as present in hypertrophic induction medium (36), *e.g.* phosphate (37). However, it remains speculative to what extent the factors in hypertrophic media induce hypertrophic differentiation *in vivo* (38), as until now these have remained elusive.

### PTH decreased DNA content of cartilaginous constructs

Since PTHrP stimulates growth plate chondrocyte proliferation (34), the inhibitory effect of PTH on the tissue constructs’ DNA content was rather surprising. However, the direction of the PTH-mediated effects appears to be dependent on dose (39,40) and maturation phase (41). The latter is stressed by the fact that PTH was able to increase the mitogenic response of growth plate chondrocytes, but not of healthy articular chondrocytes (42). The question arises which difference between growth plate and articular chondrocytes may explain this differential response to PTH. While at a gene expression level there are similarities in the spatial architecture of growth plate and articular cartilage (43), their embryonic origin is distinctly different. Articular chondrocytes are embryonically derived from the interzone cells forming both the synovial lining and the articular cartilage of the joint (44) and in postnatal stages can be distinguished from growth plate chondrocytes by lubricin expression (45). Possibly, the signaling mechanisms are also different between articular and growth plate chondrocytes. Lastly, the inhibitory effect of PTH on the DNA content could have been caused by signaling via a different receptor. For example, binding of the PTH family member TIP39 to PTHR2 has been shown to inhibit articular chondrocyte proliferation (46). Future studies should further determine the age- and possibly OA-dependent expression and effects of PTHR2 versus PTHR1 signaling in articular chondrocytes.

### PTH decreased GAG and type II collagen deposition of cartilaginous constructs

0.1 µM PTH treatment has been shown to stimulate collagen type II deposition in hypertrophic articular chondrocytes (35). Therefore, the second aim of this study was to determine whether PTH could also enhance cartilaginous (GAG- and type II collagen-rich) extracellular matrix deposition by healthy human articular chondrocytes. Mainly due to the decrease in cell number, GAG and type II collagen deposition were inhibited by 0.1 and 1 µM PTH in the current study. The latter is in line with results obtained with rat articular chondrocytes (10) and chicken sterna (47). Possibly, lowering the PTH concentration to below 0.1 µM could have prevented the observed inhibition of type II collagen deposition (48), since PTH is known to exert opposite effects at low versus high concentrations in this respect (39,40).

### Clinical application of PTH

When considering the clinical application of PTH for the repair of cartilage defects, intermittent PTH administration seems favorable to continuous administration: it enhanced chondrogenesis and suppressed hypertrophy of human MSCs compared with continuous administration (16,17), improved the quality of regenerated rabbit cartilage (21) and prevented osteoporosis in postmenopausal women (49). In the current study, PTH was administered continuously instead of intermittently, possibly explaining the absence of a beneficial effect. However, local sustained delivery resulted in similar protective effects against GAG loss and type X collagen deposition in a rat OA model as obtained with intermittent injection of PTH (50), suggesting that local delivery may overcome the requirement for intermittent exposure.

The inhibition of cartilage regeneration observed in our study would argue against the use of PTH in cartilage repair procedures based on ACI. Although PTH did not exert any positive effects on healthy human articular chondrocytes, it could still be beneficial for cartilage repair based on MSCs alone (13–17) (*e.g.* microfractures) or procedures in which an overwhelming concentration of MSCs is combined with chondrocytes, for example during a one-stage cartilage repair procedure. However, care should be taken not to interfere with endogenous repair by for example the cartilage progenitors or interzone-derived MSCs from the synovial lining, which are known to contribute to the repair of cartilage defects (44).

## Conclusions

Continuous PTH treatment inhibited regeneration of healthy human articular chondrocytes, while hypertrophic differentiation was not influenced, indicating that articular chondrocytes respond differently to PTH compared with MSCs. Therefore, PTH may be more suitable for cartilage repair based on MSCs than on articular chondrocytes.

## Acknowledgements

The authors would like to thank Marianna Tryfonidou for critically reviewing this manuscript before submission. There are no known conflicts of interest associated with this publication. LC and FB are supported by the funding from the Dutch Arthritis Foundation (LLP12 and 22, respectively). Part of the work was also funded by the Anna Foundation for Musculoskeletal Research in the Netherlands and the Netherlands Organisation for Health Research and Development (NWO). There has been no significant financial support for this work that could have influenced its outcome. Funders did not play any role in: study design; collection, analysis, or interpretation of data; manuscript writing; or the decision to submit the manuscript for publication.

## Author contributions statement

The manuscript has been read and approved by all authors and there are no other persons who satisfied the criteria for authorship. Each author has made substantial contributions to all three of sections (1) the conception and design of the study, or acquisition of data, or analysis and interpretation of data, (2) drafting the article or revising it critically for important intellectual content, (3) final approval of the version to be submitted. Given the extent of the study, 8 authors have been listed. Conception and design: LC, MR, FB. Analysis and interpretation of the data: LV, LC, FB. Drafting of the article: MR, FB. Critical revision of the article for important intellectual content: LC. Statistical expertise: MR, LC. Obtaining of funding: LC. Administrative, technical, or logistic support: VA, MR, LV. Collection and assembly of data: MR, MR, VA, FC. All authors read and approved the final manuscript. LC takes responsibility for the integrity of the work as a whole, from inception to finished article.

## References

(1) Welch T, Mandelbaum B, Tom M. Autologous Chondrocyte Implantation: Past, Present, and Future. Sports Med Arthrosc Rev 2016 Jun;24(2):85–91.

(2) Knutsen G, Engebretsen L, Ludvigsen TC, Drogset JO, Grontvedt T, Solheim E, et al. Autologous chondrocyte implantation compared with microfracture in the knee. A randomized trial. J Bone Joint Surg Am 2004 Mar;86-A(3):455–464.

(3) Zhang W, Chen J, Zhang S, Ouyang HW. Inhibitory function of parathyroid hormone-related protein on chondrocyte hypertrophy: the implication for articular cartilage repair. Arthritis Res Ther 2012 Aug 31;14(4):221.

(4) Mueller MB, Tuan RS. Functional characterization of hypertrophy in chondrogenesis of human mesenchymal stem cells. Arthritis Rheum 2008 May;58(5):1377–1388.

(5) Amizuka N, Warshawsky H, Henderson JE, Goltzman D, Karaplis AC. Parathyroid hormone-related peptide-depleted mice show abnormal epiphyseal cartilage development and altered endochondral bone formation. J Cell Biol 1994 Sep;126(6):1611–1623.

(6) de Crombrugghe B, Lefebvre V, Nakashima K. Regulatory mechanisms in the pathways of cartilage and bone formation. Curr Opin Cell Biol 2001 Dec;13(6):721–727.

(7) Zerega B, Cermelli S, Bianco P, Cancedda R, Cancedda FD. Parathyroid hormone [PTH(1-34)] and parathyroid hormone-related protein [PTHrP(1-34)] promote reversion of hypertrophic chondrocytes to a prehypertrophic proliferating phenotype and prevent terminal differentiation of osteoblast-like cells. J Bone Miner Res 1999 Aug;14(8):1281–1289.

(8) Weisser J, Riemer S, Schmidl M, Suva LJ, Poschl E, Brauer R, et al. Four distinct chondrocyte populations in the fetal bovine growth plate: highest expression levels of PTH/PTHrP receptor, Indian hedgehog, and MMP-13 in hypertrophic chondrocytes and their suppression by PTH (1-34) and PTHrP (1-40). Exp Cell Res 2002 Sep 10;279(1):1–13.

(9) Pioszak AA, Parker NR, Gardella TJ, Xu HE. Structural basis for parathyroid hormone-related protein binding to the parathyroid hormone receptor and design of conformation-selective peptides. J Biol Chem 2009 Oct 9;284(41):28382–28391.

(10) Tsukazaki T, Ohtsuru A, Namba H, Oda J, Motomura K, Osaki M, et al. Parathyroid hormone-related protein (PTHrP) action in rat articular chondrocytes: comparison of PTH(1-34), PTHrP(1-34), PTHrP(1-141), PTHrP(100-114) and antisense oligonucleotides against PTHrP. J Endocrinol 1996 Sep;150(3):359–368.

(11) Huang W, Zhou X, Lefebvre V, de Crombrugghe B. Phosphorylation of SOX9 by cyclic AMP-dependent protein kinase A enhances SOX9’s ability to transactivate a Col2a1 chondrocyte-specific enhancer. Mol Cell Biol 2000 Jun;20(11):4149–4158.

(12) Zhang M, Xie R, Hou W, Wang B, Shen R, Wang X, et al. PTHrP prevents chondrocyte premature hypertrophy by inducing cyclin-D1-dependent Runx2 and Runx3 phosphorylation, ubiquitylation and proteasomal degradation. J Cell Sci 2009 May 1;122(Pt 9):1382–1389.

(13) Kafienah W, Mistry S, Dickinson SC, Sims TJ, Learmonth I, Hollander AP. Three-dimensional cartilage tissue engineering using adult stem cells from osteoarthritis patients. Arthritis Rheum 2007 Jan;56(1):177–187.

(14) Kim YJ, Kim HJ, Im GI. PTHrP promotes chondrogenesis and suppresses hypertrophy from both bone marrow-derived and adipose tissue-derived MSCs. Biochem Biophys Res Commun 2008 Aug 15;373(1):104–108.

(15) Mwale F, Yao G, Ouellet JA, Petit A, Antoniou J. Effect of parathyroid hormone on type X and type II collagen expression in mesenchymal stem cells from osteoarthritic patients. Tissue Eng Part A 2010 Nov;16(11):3449–3455.

(16) Fischer J, Aulmann A, Dexheimer V, Grossner T, Richter W. Intermittent PTHrP(1-34) exposure augments chondrogenesis and reduces hypertrophy of mesenchymal stromal cells. Stem Cells Dev 2014 Oct 15;23(20):2513–2523.

(17) Fischer J, Ortel M, Hagmann S, Hoeflich A, Richter W. Role of PTHrP(1-34) Pulse Frequency Versus Pulse Duration to Enhance Mesenchymal Stromal Cell Chondrogenesis. J Cell Physiol 2016 Dec;231(12):2673–2681.

(18) Bach FC, Rutten K, Hendriks K, Riemers FM, Cornelissen P, de Bruin A, et al. The paracrine feedback loop between vitamin D(3) (1,25(OH)(2)D(3)) and PTHrP in prehypertrophic chondrocytes. J Cell Physiol 2014 Dec;229(12):1999–2014.

(19) Iwamoto M, Jikko A, Murakami H, Shimazu A, Nakashima K, Iwamoto M, et al. Changes in parathyroid hormone receptors during chondrocyte cytodifferentiation. J Biol Chem 1994 Jun 24;269(25):17245–17251.

(20) Kudo S, Mizuta H, Otsuka Y, Takagi K, Hiraki Y. Inhibition of chondrogenesis by parathyroid hormone in vivo during repair of full-thickness defects of articular cartilage. J Bone Miner Res 2000 Feb;15(2):253–260.

(21) Kudo S, Mizuta H, Takagi K, Hiraki Y. Cartilaginous repair of full-thickness articular cartilage defects is induced by the intermittent activation of PTH/PTHrP signaling. Osteoarthritis Cartilage 2011 Jul;19(7):886–894.

(22) de Windt TS, Vonk LA, Slaper-Cortenbach ICM, Nizak R, van Rijen MHP, Saris DBF. Allogeneic MSCs and Recycled Autologous Chondrons Mixed in a One-Stage Cartilage Cell Transplantion: A First-in-Man Trial in 35 Patients. Stem Cells 2017 Aug;35(8):1984–1993.

(23) Cournil-Henrionnet C, Huselstein C, Wang Y, Galois L, Mainard D, Decot V, et al. Phenotypic analysis of cell surface markers and gene expression of human mesenchymal stem cells and chondrocytes during monolayer expansion. Biorheology 2008;45(3-4):513–526.

(24) Benz K, Stippich C, Freudigmann C, Mollenhauer JA, Aicher WK. Maintenance of “stem cell” features of cartilage cell sub-populations during in vitro propagation. J Transl Med 2013 Jan 30;11:27-5876-11-27.

(25) Collins DH. The Pathology of Articular and Spinal Diseases. London: Arnold; 1949.

(26) van Diest PJ. No consent should be needed for using leftover body material for scientific purposes. BMJ 2002;325:648.

(27) Gao L, Orth P, Cucchiarini M, Madry H. Autologous Matrix-Induced Chondrogenesis: A Systematic Review of the Clinical Evidence. Am J Sports Med 2017 Nov 1:363546517740575.

(28) Kandel RA, Chen H, Clark J, Renlund R. Transplantation of cartilagenous tissue generated in vitro into articular joint defects. Artif Cells Blood Substit Immobil Biotechnol 1995;23(5):565–577.

(29) Rutgers M, Saris DB, Vonk LA, van Rijen MH, Akrum V, Langeveld D, et al. Effect of collagen type I or type II on chondrogenesis by cultured human articular chondrocytes. Tissue Eng Part A 2013 Jan;19(1-2):59–65.

(30) Rosenberg L. Chemical basis for the histological use of safranin O in the study of articular cartilage.. J Bone Joint Surg Am 1971;53:69–82.

(31) Grogan SP, Barbero A, Winkelmann V, Rieser F, Fitzsimmons JS, O’Driscoll S, et al. Visual histological grading system for the evaluation of in vitro-generated neocartilage. Tissue Eng 2006 Aug;12(8):2141–2149.

(32) Whiteman P. The quantitative measurement of Alcian Blue-glycosaminoglycan complexes. Biochem J 1973 Feb;131(2):343–350.

(33) Kim YJ, Sah RL, Doong JY, Grodzinsky AJ. Fluorometric assay of DNA in cartilage explants using Hoechst 33258. Anal Biochem 1988 Oct;174(1):168–176.

(34) Kronenberg HM. Developmental regulation of the growth plate. Nature 2003 May 15;423(6937):332–6.

(35) Chang LH, Wu SC, Chen CH, Wang GJ, Chang JK, Ho ML. Parathyroid hormone 1-34 reduces dexamethasone-induced terminal differentiation in human articular chondrocytes. Toxicology 2016 Aug 10;368–369:116-128.

(36) Gawlitta D, van Rijen MH, Schrijver EJ, Alblas J, Dhert WJ. Hypoxia impedes hypertrophic chondrogenesis of human multipotent stromal cells. Tissue Eng Part A 2012 Oct;18(19-20):1957–1966.

(37) Bellows CG, Heersche JN, Aubin JE. Inorganic phosphate added exogenously or released from beta-glycerophosphate initiates mineralization of osteoid nodules in vitro. Bone Miner 1992 Apr;17(1):15–29.

(38) Zhang R, Lu Y, Ye L, Yuan B, Yu S, Qin C, et al. Unique roles of phosphorus in endochondral bone formation and osteocyte maturation. J Bone Miner Res 2011 May;26(5):1047–1056.

(39) Zhang Y, Kumagai K, Saito T. Effect of parathyroid hormone on early chondrogenic differentiation from mesenchymal stem cells. J Orthop Surg Res 2014 Aug 1;9:68-014-0068-5.

(40) Weiss S, Hennig T, Bock R, Steck E, Richter W. Impact of growth factors and PTHrP on early and late chondrogenic differentiation of human mesenchymal stem cells. J Cell Physiol 2010 Apr;223(1):84–93.

(41) Lossdorfer S, Gotz W, Rath-Deschner B, Jager A. Parathyroid hormone(1-34) mediates proliferative and apoptotic signaling in human periodontal ligament cells in vitro via protein kinase C-dependent and protein kinase A-dependent pathways. Cell Tissue Res 2006 Sep;325(3):469–479.

(42) Crabb ID, O’Keefe RJ, Puzas JE, Rosier RN. Differential effects of parathyroid hormone on chick growth plate and articular chondrocytes. Calcif Tissue Int 1992 Jan;50(1):61–66.

(43) Chau M, Lui JC, Landman EB, Spath SS, Vortkamp A, Baron J, et al. Gene expression profiling reveals similarities between the spatial architectures of postnatal articular and growth plate cartilage. PLoS One 2014 Jul 28;9(7):e103061.

(44) Roelofs AJ, Zupan J, Riemen AHK, Kania K, Ansboro S, White N, et al. Joint morphogenetic cells in the adult mammalian synovium. Nat Commun 2017 May 16;8:15040.

(45) Rhee DK, Marcelino J, Baker M, Gong Y, Smits P, Lefebvre V, et al. The secreted glycoprotein lubricin protects cartilage surfaces and inhibits synovial cell overgrowth. J Clin Invest 2005 Mar;115(3):622–631.

(46) Panda D, Goltzman D, Juppner H, Karaplis AC. TIP39/parathyroid hormone type 2 receptor signaling is a potent inhibitor of chondrocyte proliferation and differentiation. Am J Physiol Endocrinol Metab 2009 Nov;297(5):E1125–36.

(47) Harrington EK, Coon DJ, Kern MF, Svoboda KK. PTH stimulated growth and decreased Col-X deposition are phosphotidylinositol-3,4,5 triphosphate kinase and mitogen activating protein kinase dependent in avian sterna. Anat Rec (Hoboken) 2010 Feb;293(2):225–234.

(48) Mueller MB, Fischer M, Zellner J, Berner A, Dienstknecht T, Kujat R, et al. Effect of parathyroid hormone-related protein in an in vitro hypertrophy model for mesenchymal stem cell chondrogenesis. Int Orthop 2013 May;37(5):945–951.

(49) Zanchetta JR, Bogado CE, Cisari C, Aslanidis S, Greisen H, Fox J, et al. Treatment of postmenopausal women with osteoporosis with PTH(1-84) for 36 months: treatment extension study. Curr Med Res Opin 2010 Nov;26(11):2627–2633.

(50) Eswaramoorthy R, Chang CC, Wu SC, Wang GJ, Chang JK, Ho ML. Sustained release of PTH(1-34) from PLGA microspheres suppresses osteoarthritis progression in rats. Acta Biomater 2012 Jul;8(6):2254–2262.

